# An integrated method for single cell proteomics with simultaneous measurements of intracellular drug concentration implicates new mechanisms for adaptation to KRAS^G12D^ inhibitors

**DOI:** 10.1101/2023.11.18.567669

**Authors:** Benjamin C. Orsburn

## Abstract

It is well established that a population of single human cells will often respond to the same drug treatment in a heterogeneous manner. In the context of chemotherapeutics, these diverse responses may lead to individual adaptation mechanisms and ultimately multiple distinct methods of resistance. The obvious question from a pharmacology perspective is how intracellular concentrations of active drug varies between individual cells, and what role does that variation play in drug response heterogeneity? To date, no integrated methods for rapidly measuring intracellular drug levels while simultaneously measuring drug responses have been described. This study describes a method for single cell preparation that allows proteins to be extracted and digested from single cells while maintaining conditions for small molecules to be simultaneously measured. The method as described allows up to 40 cells to be analyzed per instrument per day. When applied to a KRAS^G12D^ small molecule inhibitor I observe a wide degree of intracellular levels of the drug, and that proteomic responses largely stratify based on the concentration of drug within each single cell. Further work is in progress to develop and standardize this method and – more importantly – to normalize drug measurements against direct measurements of cell volume. However, these preliminary results appear promising for the identification of single cells with unique drug response mechanisms. All data described in this study has been made publicly available through the ProteomeXchange consortium under accession PXD046002.

**Abstract graphic:** 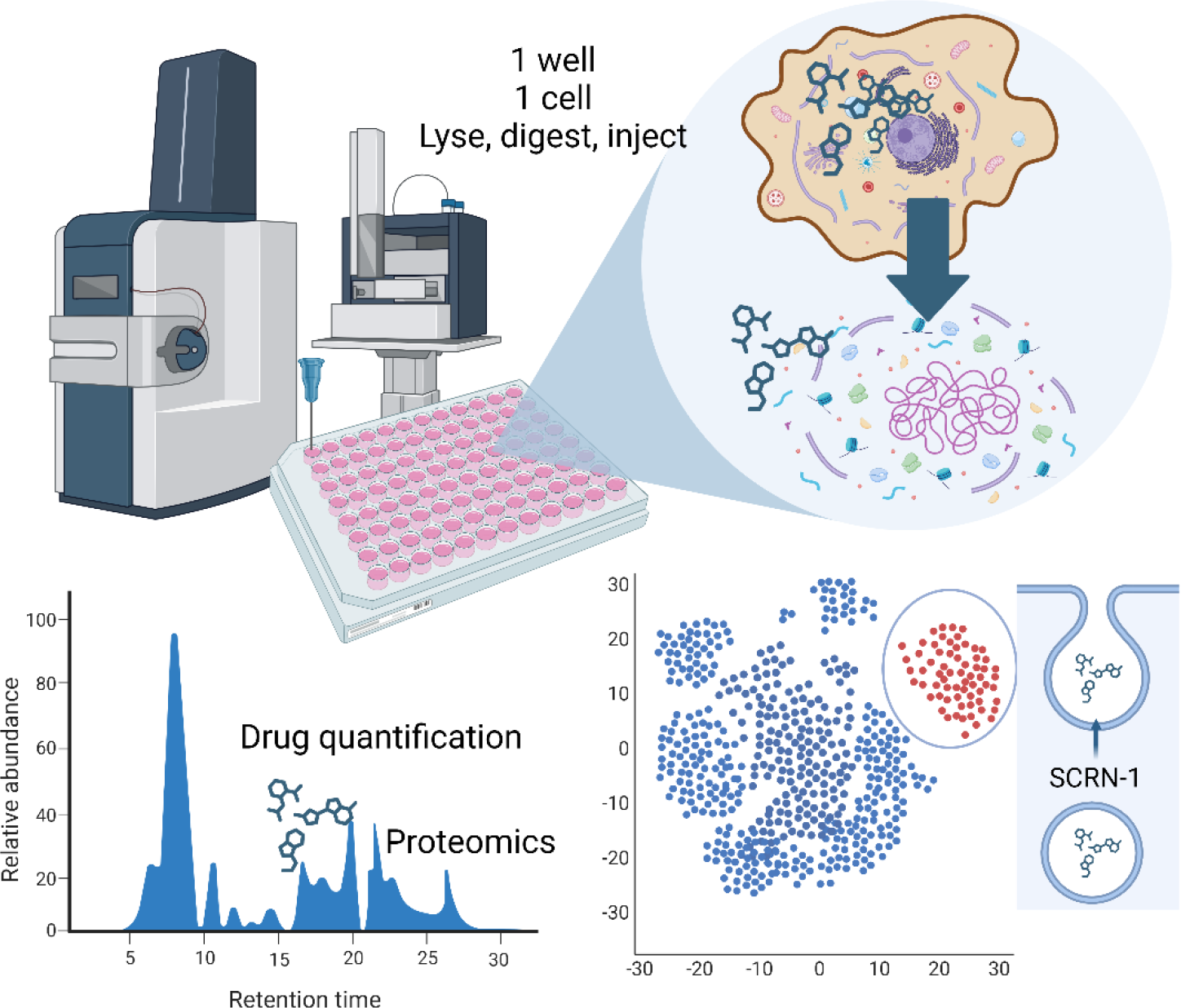

## Introduction

It is now well accepted that a population of cells will not respond to the same dose of drug in a completely uniform and homogenous manner. ^1,2^ From a basic evolutionary perspective it simply doesn’t make sense for a population of cells to exhibit uniform responses that may result in uniform destruction under any single condition. ^3^ Even clonal populations of cells that have been synchronized to the same cell cycle conditions are not truly uniform, as the release back into the normal cell cycle does not occur at identical rates. ^4^ Through techniques such as Raman spectroscopy, ^5^ inductively coupled plasma single cell mass spectrometry ^6^ and secondary ion mass spectrometry, ^7^ it is now possible to observe drug concentrations within individual cells. However, no method has described the capability to simultaneously measure intracellular drug concentrations while obtaining unbiased global information on cell state at high relative throughput.

Recent advances in sample handling, chromatography and mass spectrometry have ushered in an exciting new era and the proteomic analysis of single human cells. ^8–10^ While this field is clearly in an early growth phase with improvements coming in daily from new innovations, new tools are trickling down to all branches of mass spectrometry. Due to a new generation of low flow chromatography systems and high resolution mass spectrometers with increasing speed and sensitivity, it is fair to say that most of the challenges are due to sample loss during handling. ^11,12^ Effective methods to combat sample loss include isolating single cells and performing all steps of cell lysis and digestion in the smallest possible volume with the lowest amount of sample contact possible. To further mitigate sample loss the well or drop where the cell was deposited and processed can be directly loaded into the LCMS system. ^13,14^

Work in our group has centered on attempting to understand how the heterogeneity in response to KRAS small molecule inhibitors relates to the development of cellular resistance to these drugs. ^15^ A common question while describing these results has been whether this is simply an indirect measurement of the amount of drug that is present in each single cell. I hypothesized that with a drug with a convenient mass-to-charge ratio it would be possible to simultaneously measure intracellular drug concentrations while obtaining quantitative proteomic data from isolated single cells of a typical size. When applying this method to the KRAS^G12D^ inhibitor MRTX1133, ^16,17^ I observe a large variation in intracellular drug accumulation. Furthermore, distinct proteomic alterations correspond directly to intracellular levels of the drug and seemingly provide new insight into how cells respond and adapt to KRAS inhibition.

## Methods

### MRTX1133 Dilution curves

MRTX1133 was purchased from Cayman Chemical (Catalog 36413) and was first resuspended in 100% LCMS grade DMSO (Pierce) in a sterile cell culture hood. Aliquots were taken for resuspension for standard curves and additional aliquots were made for dosing cells. For standard curves, 10x serial dilutions were made in 0.1% formic acid 0.1% n-Dodecyl-beta-Maltoside Detergent (DDM, Thermo Fisher, 89902) in LCMS grade water. DDM was used to help replicate single cell analysis conditions. Dilutions down to an estimated 5 attogram / μL MRTX1133 were prepared in 10 μL aliquots in sealed 96 well plates (Fisher, 60180-M143) and centrifuged prior to loading on the autosampler. Dilutions curves were analyzed using an EasyNLC 1200 nanoflow chromatography system (Proxeon) coupled to a TIMSTOF SCP mass spectrometer (Bruker Daltronic). The initial proof of concept curves were analyzed on the same LC column (IonOpticks, “Elite” with integrated CaptiveSpray emitter) as single cell extracts. The IonOpticks “Elite” is a 15 cm column with 75 μm internal diameter. The final curves were analyzed using a similar 15 cm x 75 μm PepSep C-18 column with a 10 μm CaptiveSpray emitter connected by a “zero dead volume union” (both, Bruker Daltronic). The repeat dilution curves were performed with this column due to a temporary internal purchasing issue with Australian vendors. Two microliter injections of each dilution were performed in triplicate before a water blank and moving to the next higher concentration sample.

### LCMS instrument parameters

An EasyNLC 1200 system (Proxeon) coupled to a TIMSTOF SCP (Bruker Daltronic) was used for all analyses. Peptides were separated using a gradient with a constant flow of 300 nL/min with integrated CaptiveSpray emitter held at 1500V. For standard injections, 2 μL of each well was loaded with a partial loop injection of 7 μL at 900 bar. For single cell injections 4 μL was picked up with a partial loop injection volume of 10 μL to counter for peptide diffusion within the sample loop. Buffer A consisted of LCMS grade water in 0.5% acetic acid. Buffer B consisted of 80% LCMS grade acetonitrile in water acidified with the same. The 30 minute gradient used in all experiments began at 8% buffer B and ramped to 35% B by 22 minutes. The gradient then increased to 100% B by 26 minutes where it held for 2 minutes before returning to baseline conditions. The column was equilibrated in 12 microliters of baseline conditions prior to each injection.

The TIMSTOF SCP system was operated in diaPASEF mode using a method with 50 Da isolation windows provided by Dr. Michael Krawitzky of Bruker Daltronic. In this method, the MS1 scan is obtained from 100-1700 m/z with a 1/k0 window of 0.7 – 1.4. Ions that enter the mass analyzer in these systems are restricted by a user defined polygon that imparts dramatic alterations on ion signal measured and must be carefully optimized for each series of experiments. In this case, the polygon began at 295 m/z and attempted to not acquire any ions in the MS1 cloud below 595 m/z. The method utilizes 6 cycles with 3 isolation windows per cycle resulting in a total time of 0.72 seconds with 100 ms ramp time. The “high sensitivity” detector mode was used in all instances.

### Isolation and preparation of PANC0203 cancer cells

The PANC 0203 cancer cells used in this study were obtained from ATCC and grown following vendor instructions with RPMI 1640 (ATCC 30-2001) supplemented with 15% fetal bovine serum (ATCC 30-2020). Culture media was further supplemented with 10 units of human insulin (Fisher) and 10 mg/mL Penn Strep antibiotic solution (ATCC 30-2300). All cell lines were passaged a minimum of 3 times prior to single cell isolation. A summary of the cell harvest method is illustrated in **Figure 1**. Cells were harvested first by vacuum aspiration of the cell culture media. The adherent cells were briefly rinsed in 3 mL of 0.05% Trypsin plus EDTA solution (ATCC 30-2001). This solution was rapidly aspirated off and replaced with 3 mL of the same solution. The cells were examined by light field microscopy and incubated at 37°C with multiple examinations until the adherent cells had lifted off the plate surface. The active trypsin was then quenched by the addition of 7 mL of the original culture media. The 10 mL solution was transferred to sterile 15 mL Falcon tubes (Fisher) and centrifuged at 300 *x g* for 3 minutes to pellet the cells. The supernatant was gently aspirated off and the cells were resuspended in PBS solution without calcium or magnesium with 0.1% BSA (both, Fisher Scientific) at 1 million cells per mL as estimated by bright field microscopy. Cells for single cell aliquoting were gently dissociated from clumps by slowly pipetting a solution of approximately 1 million cells through a Falcon cell strainer (Fisher, 353420) and the cells were placed on wet ice and immediately transported to the JHU Public Health sorting core. Non-viable cells were labeled with a propidium iodide (PI) solution provided by the core facility and briefly vortexed prior to cell isolation and aliquoting. The PI solution was the only stain utilized and was used to isolate cells with ruptured membranes and exclude them from isolation and deposition.

**Figure 1.**
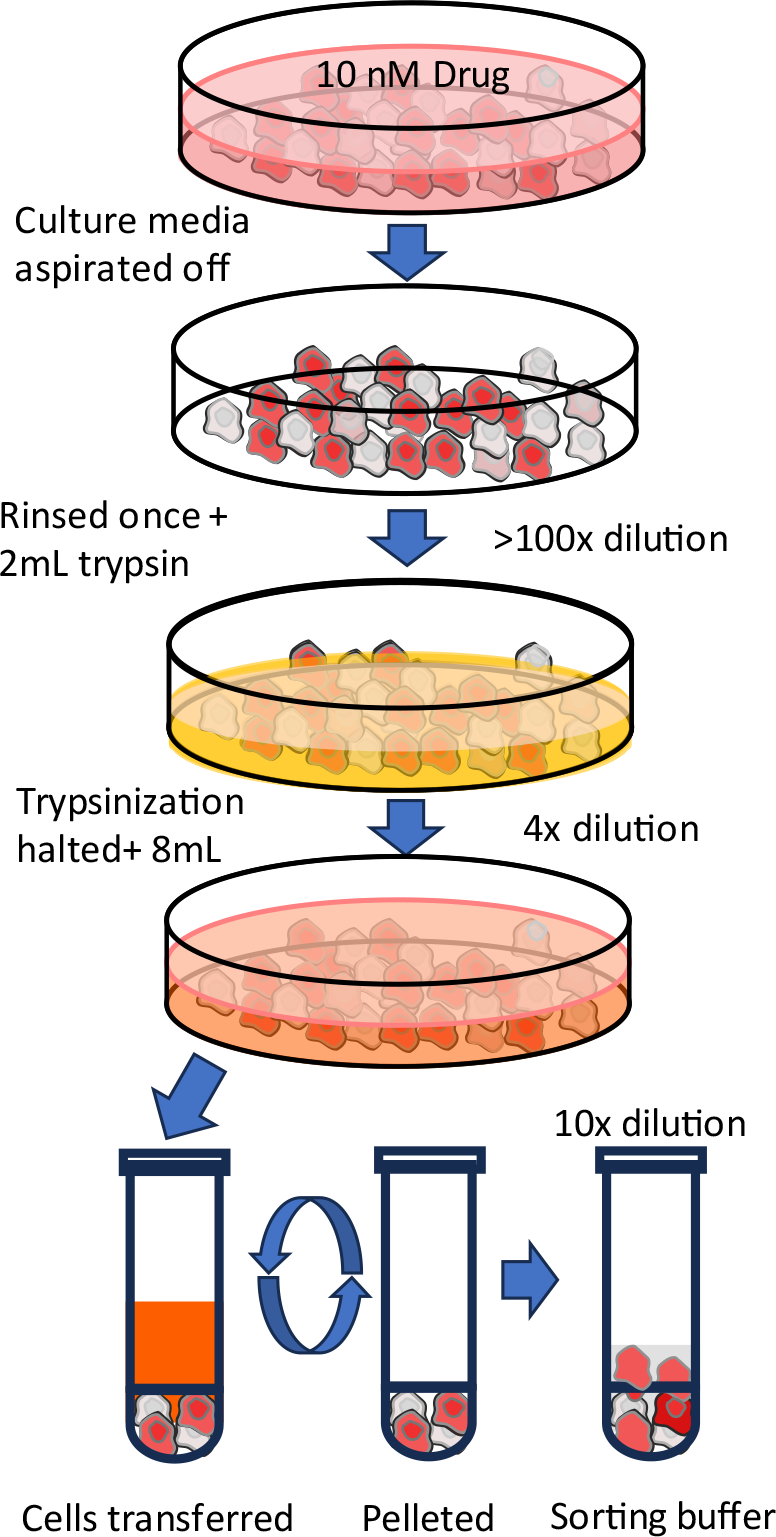
An overview of the harvesting and preparation of drug treated cells for single cell aliquoting.

**Figure 2.**
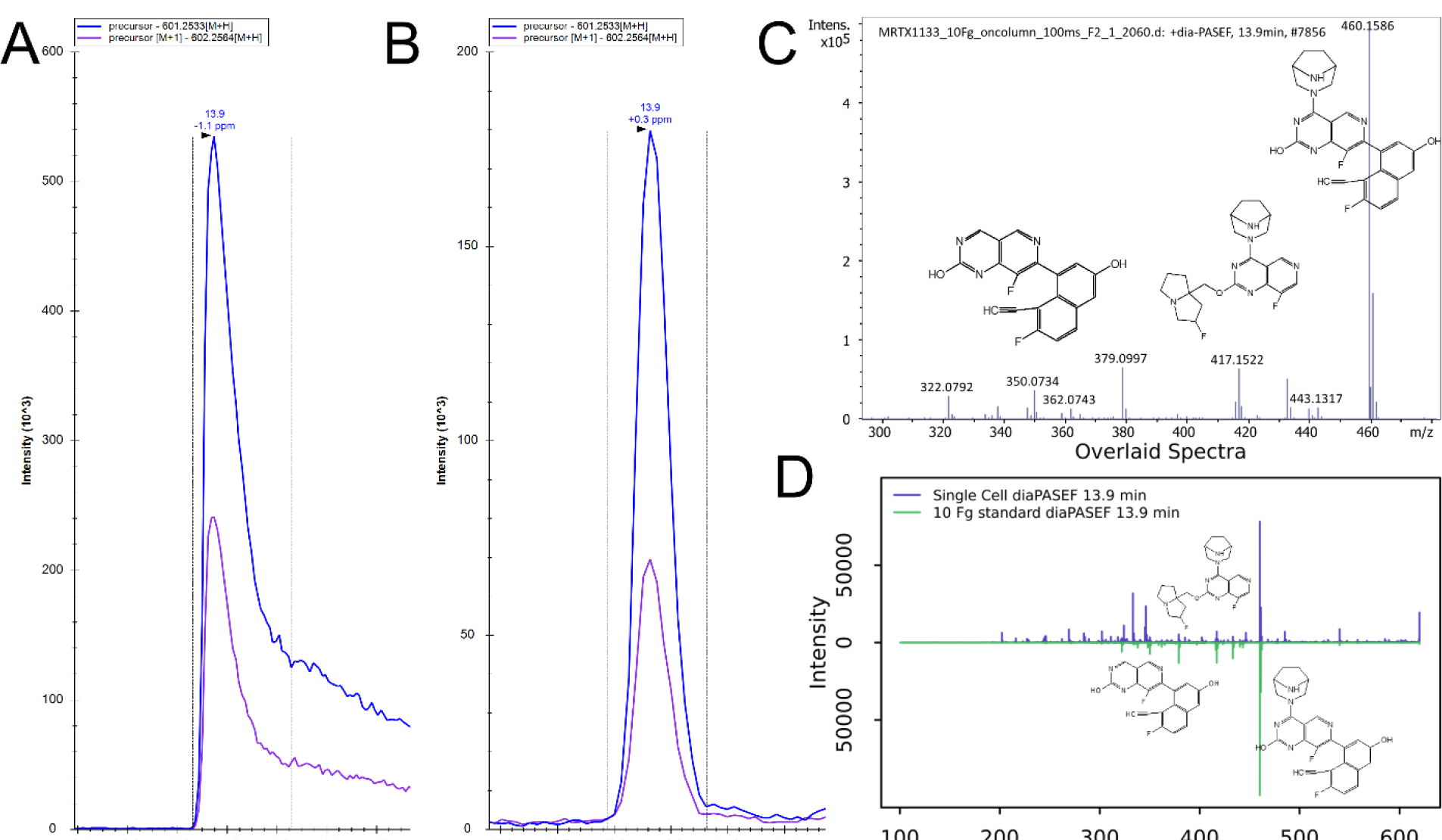
MRTX1133 diluted standard and detection in single cells. **A**. The upper limit of quantitation of MRTX1133 at 10 femtogram on column exhibiting peak broadening on this system. **B**. Two isotopes of MRTX1133 measured in a single drug treated PANC 02.03 cell. **C**. Fragmentation pattern of the drug standard. **D**. Mirror plots of diaPASEF scan from a single PANC 02.03 cell at 13.9 minutes (top panel) and from the MRTX1333 standard at 10 femtogram at 13.9 minutes.

Single cells were aliquoted using an analog MoFlo sorter into cold 96 well plates containing 2 microliters of LCMS grade acetonitrile. At the completion of each plate aliquoting they were immediately sealed and placed in an insulated box of dry ice with the wells pressed into the material to ensure rapid cooling. The frozen single cells were transported back to -70°C storage. For processing, acetonitrile was driven off by heating the cells for 90 seconds on a hotplate at 95°C. Dried cell lysate was digested using a solution of 5 nanogram/microliter LCMS grade trypsin (Pierce) in 0.1% n-Dodecyl-beta-Maltoside Detergent (DDM, Thermo Fisher, 89902) and 50mM TEAB. Two microliters of trypsin solution were used for each cell prior to the plate being tightly sealed with adhesive plate tape (Fisher, 60180-M143) and room temperature overnight digestion. Following digestion, the peptide digest was briefly centrifuged at 4°C to condense evaporation and the plates were completely dried with vacuum centrifugation. The peptides were resuspended in 3.5 μL of 0.1% formic acid, vortexed in tightly sealed plates and centrifuged prior to loading on the autosampler.

## Data Analysis

SpectroNaut 18 (Biognosys) was used for all proteomic data analysis using the directDIA+ workflow and a human Uniprot library utilizing all default parameters for data calibration and analysis. All files were first converted to HT using the appropriately named “HT onverter” so tware rom Biognosys prior to analysis. MRTX1133 concentrations were calculated using Skyline 21.2.0.568. ^18^ Due to the fact the author doesn’t know how to per orm a standard curve calculation in this so tware and he suspects that no one else does either, the area under the curve for each peak was extracted and the curves were plotted in Excel 365 and GraphPad Prizm 10.0.1. For analysis of single cell data, our in house software package SCP-Viz ^19^ was used to identify and extract single cells with specific characteristics, such as higher or lower relative Histone H4 concentrations. For statistical analysis of the proteomic data, the quantitative proteomic data was uploaded into the SimpliFi Cloud interpretation interface (ProtiFi, LLC) following clustering along the subpopulations identified by SCP-Viz.

## Data Availability Statement

All Bruker .d files, SpectroNaut processed results and Skyline files have been deposited at the ProteomeXchange MASSIVE partner repository and can be accessed as accession PXD046002.

## Results

### A KRASG12D inhibitor can be detected at attogram concentrations

A serial dilution of MRTX1133 could be detected using a diaPASEF proteomics method optimized to allow the limited fragmentation of lower mass ions. An estimated limit of detection of 100 attograms on column (0.164 attomole) was obtained. The upper limit of quantification appears to be approximately 10 femtograms on column (16.64 attomole) due to peak broadening on the IonOpticks chromatography column, although the total response of the area under the curve appears to maintain linearity at this level.

### The TIMS exclusion polygon can be optimized to collect MRTX1133 ions and tryptic peptides

Trapped ion mobility mass spectrometry has proven a disruptive technology in proteomics, imaging mass spectrometry, metabolomics and lipidomics. In a traditional proteomic workflow on a TIMSTOF platform, a visual tool is used to select a mass and ion mobility window for analysis. Due to the fact that most tryptic peptides have at least two positive charges at LCMS buffer conditions, singly charged ions are placed outside of these cutoff windows. By carefully drawing this exclusion polygon to accept singly charged ions down to an m/z of 595, the MRTX1133 ion can be accumulated for MS1 and MS/MS fragmentation while tryptic peptides are also selected. While this likely has detrimental effects on total proteomic coverage, up to 1,240 proteins were identified in a single cell, with a mean of approximately 450 proteins. Protein identifications were made using a library free approach with data searched against the entire annotated human proteome. Multiple studies have shown that the number of proteins identified per single cell is lowest when using this approach alone and that SCP specific informatics and custom libraries will improve total protein numbers. ^20–23^ A more in depth analysis will be performed on a larger cohort of cells in work as described later in this manuscript.

### MRTX1133 can be confirmed in single cell isolates with multiple dimensions of confidence

In a recent metabolomic analysis of four cancer cell lines treated with MRTX1133 I observed that the drug ionized with high efficiency and can be readily detected in positive ionization mode LCMS using reversed phase chromatography. ^24^ Similarly, MRTX1133 could be readily detected in nearly all isolated drug treated single cells following sample processing. Two isotopes of MRTX1133 could be matched in serial dilutions (**Figure 1A**) and in single cells (**Figure 1B**) within tight retention time windows. In addition, the fragmentation of MRTX1133 using this standard diaPASEF method results in a dominant fragment ion with an m/z of 460.15. Additional fragment ions at 417.15 and 350.07 m/z, respectively, can be readily attributed to the subsequent loss of each functional group from the parent molecule (**Figure 1C**). All three of these ions can be distinguished in the same diaPASEF isolation windows in PANC 02.03 cells (**Figure 1D**) within strict retention time tolerance windows. In no case was MRTX1133 identified with intensity greater than the estimated limited of detection of 100 attograms on column in control PANC 02.03 single cells or in method blank control wells (**Figure 3**).

**Figure 3.**
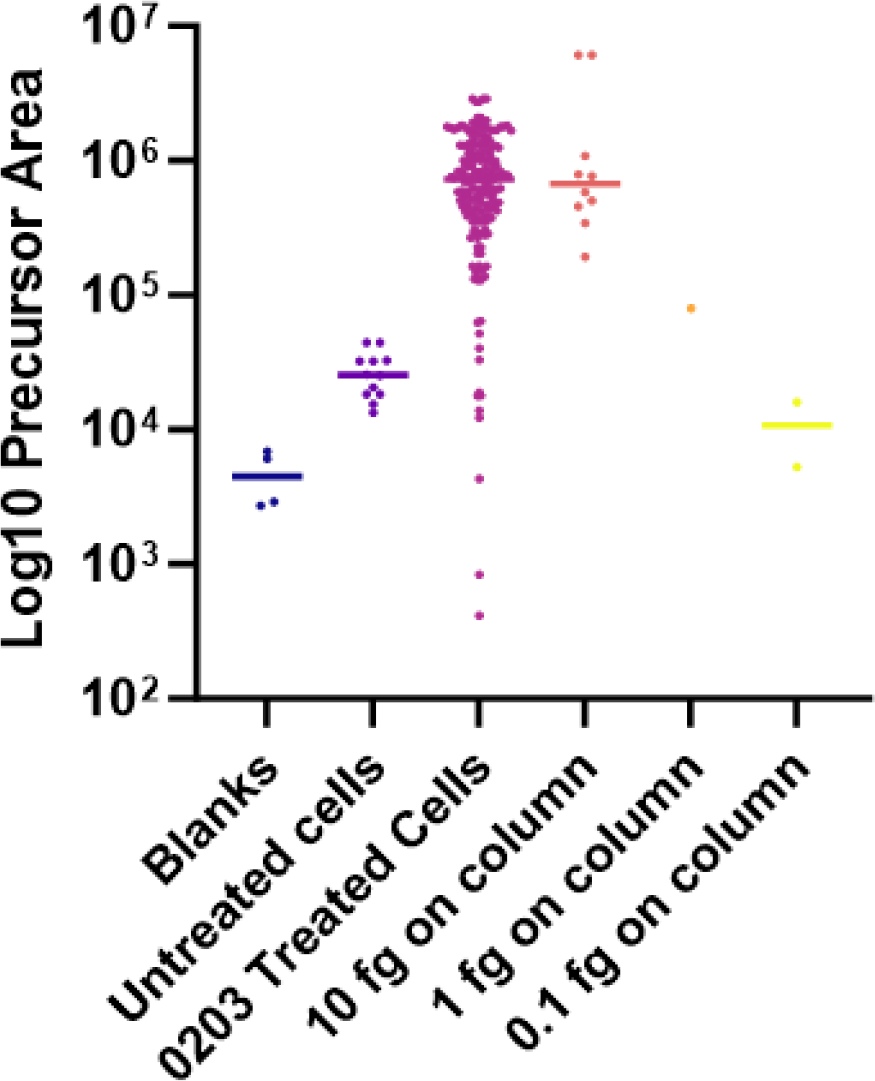
The distribution of observed MRTX1133 concentrations in single cell extracts with the median represented as well as the value for each replicate or single cell.

### MRTX1133 must be intracellular at the point the cells are sorted to be detectable

PANC 02.03 is an adherent cell line which must be trypsinized to be released from the plate surface. This process requires that the cell culture media is first removed, then rinsed with a trypsin solution, which is also aspirated off. While residual media remains after these aspirations, these are further diluted by the addition of the trypsin for the incubation step. To inactivate the trypsin, it is then diluted in cell culture media for pelleting of the cells. The diluted trypsin/media solution is again aspirated off before the cells are resuspended in cell sorting buffer. **Figure 1** is an illustration of the sample preparation process with estimates of the minimum dilution that would occur at each stage of the process. The final result is that residual MRTX1133 within the cell culture media would be diluted orders of magnitude below the current estimate of the limits of detection. Drug levels detected can only be due to storage within the intact cellular membrane at the point of single cell isolation.

### MRTX1133 levels in single cells do not correspond to indirect measurements of cell size

Single cells can vary dramatically in size due to different growth conditions and depending on their relative cell cycle stage. Recent work concluded that relative levels of histone H4 levels within a single cell can aid in estimating the relative internal volume of each single cells. ^25^ The majority of other work in single cell proteomics appears to leverage the uncorrected total ion current of the peptides observed in each single cell as an proxy for cell size for normalization purposes. ^26–28^ Surprisingly the cells with the highest relative ion current and histone H4 levels were not the cells with the highest relative amounts of MRTX1133 (**Figure 4A**).

**Figure 4.**
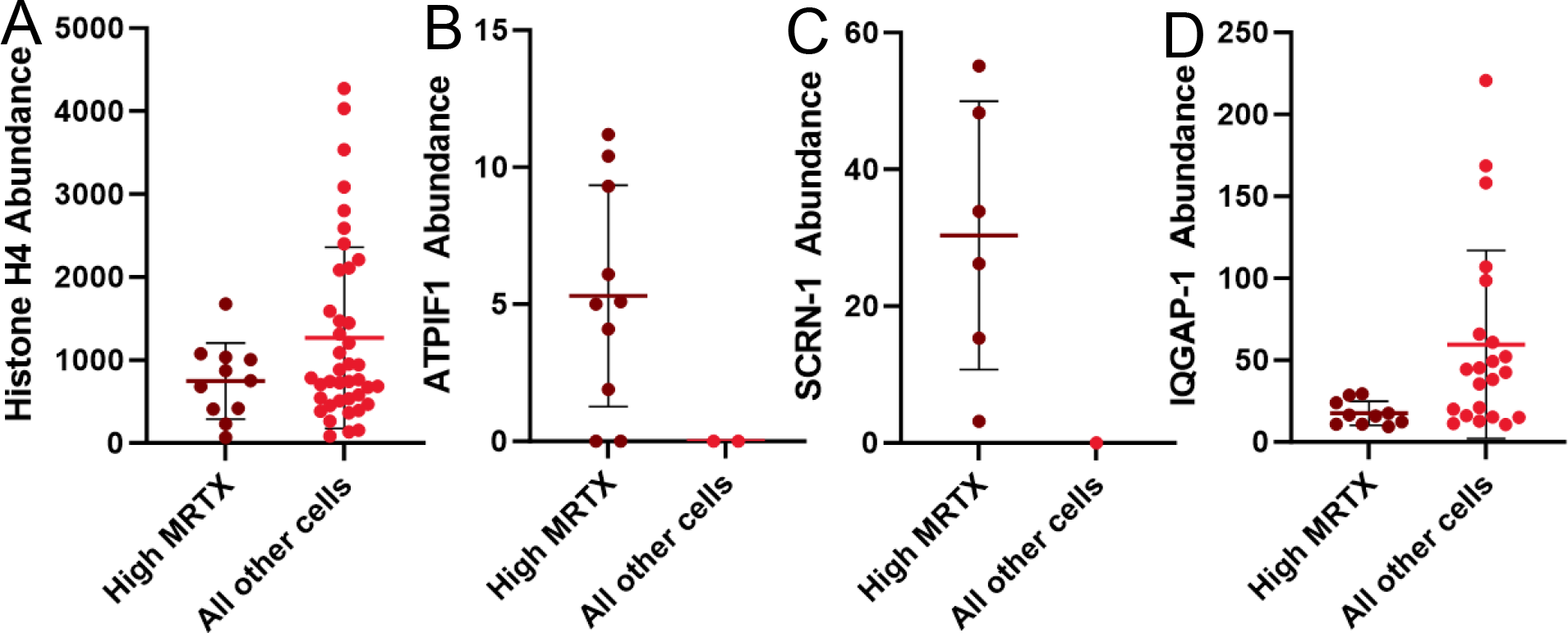
Protein expression patterns of cells striated by high MRTX1133 abundance with mean and standard deviation overlaid on single cell measurements. **A**. Histone H4 levels **B**. Mitochondrial ATP synthase inhibitor ATPI51 which is exclusively found in cells with the highest drug levels. **C**. Secernin-1 which appears follows a similar pattern to ATPIF1. **D**. The RAS interacting protein IQGAP-1 which appears repressed in cells with higher MRTX1133 levels.

### Distinct proteomic alterations are observed in single cells with higher observed drug concentration

To further evaluate proteome specific alterations consistent with higher relative intracellular concentration, I first performed a simple correlation analysis between the 210 proteins at were detected in at least 50% of all cells versus the observed MRTX1133 concentrations. The only protein observed with a positive correlation with MRTX1133 was Dermicidin (P81605) with an R^2^ of 0.4 at a p value just below 0.05. Weak negative, though statistically significant negative correlations were observed in Myoferlin, Stress-70, SLC3A2 and Galectin-3 at between -0.37 and -0.41. Due to the low strength of these correlations and the larger cohort of cells to be analyzed in follow-up work, I chose not to follow up on these proteins at this time.

To further evaluate if stratification of phenotype occurs at different MRTX1133 levels, I relabeled cells within pectroNaut ased on high, medium or low relative TX 33 levels with “high” eing de ined as more than 1 standard deviation above the mean of 135 cells where the drug was quantified above the previously stated limit of detection. “ow” was used or cells with o served TX 33 levels more than 1 standard deviation below the mean. Upon reclustering, the output data for the low, medium and high cells were imported into ProtiFi for statistical analysis of the proteomic differences between each set of cells. The complete data output for 54 cells can be accessed online through the interactive SimpliFi web portal at this open link: https://tinyurl.com/SCPplusDrug. Although this is a small set of single cells, there is a remarkable level of stratification in proteins observed in the cells with the highest drug concentrations. The ATP synthase inhibitory factor ATPIF1 (**Figure 4B**) and the exocytosis protein SCRN-1 (**Figure 4C**) are exclusively observed, and with relatively high abundance, in cells with the highest observed MRTX1133 levels. Conversely, the RAS binding partner IQGAP-1 is identified in nearly all cells in this study and is lower in nearly every case in the cells with the highest MRTX1133 concentrations (**Figure 4D**). As ATP depletion was previously observed in MRTX1133 drug treated cells, the expression of an ATPase inhibitor in a subpopulation of cells appears an interesting and contradictory observation. ^24^ While limited work has been performed on SCRN-1, a previous study demonstrated that EGFR inhibitors, which directly interact with the RAS pathway, can be potentiated by lowering SCRN-1 protein expression. ^29^ Taken together, however, it is tempting to think that a subpopulation of cells might utilize a SCRN-1 mediated exocytosis adaptation mechanism to trap KRAS proteins that have been inhibited by MRTX1133. Finally, while the pathways KRAS mutant proteins participate in are still not fully elucidated, subpopulations with altered expression of one of the main high abundance interactors of RAS proteins ^30^ suggests a great deal of variation exists between these two populations at the molecular level.

### Limitations of current study

This proof of concept work, as described has obvious and overarching limitations that must be addressed to move this study forward. The two clearest limitations can be approached in a single approach using hardware available on campus that no one currently in our group has mastered, due primarily to the large number of graduations our group has had recently. Both the low number of cells analyzed and the lack of direct measurements of cell size should be solved by employing recently described workflows on the CellenOne cell isolation and processing system. ^23^ This device has been demonstrated to obtain high accuracy measurements of cell size during cell isolation, ^31^ and to dramatically improve material recovery in single cells. ^11,22^ Furthermore, while SCRN-1 and other proteins are implicated in this study, functional studies will be necessary to support these observations. The main goal of this preprint is to attract the interest of potential collaborators with the ability to help generate siRNA or other inhibitors to further evaluate the effects these proteins have on adaptation to this drug.

## Conclusions

Around the limitations as described there is reason for optimism in this approach. First, this is a relatively simple sample preparation workflow that yields proteomics data and reasonable estimates of internal drug concentrations. Although the compromises necessary for the ion mobility isolation led to reductions in protein identifications, multiple studies have demonstrated that TIMS windows can be altered beyond the capabilities of the vendor provided software. Due to the amount of development in the diaPASEF space with new approaches like SLICE-PASEF, ^32^ pyDIAID, ^33^ and miDIApasef ^34^ the next generation of TIMS Control software features more advanced capabilities for TIMS isolation than our current software version. With superior control over the ions selected for isolation and fragmentation, I anticipate that drugs with relatively convenient m/z and ion mobilities will soon be quantifiable with less loss in proteins identified per cell. In addition, the preparation of cells in 96-well plates leads to enormous losses in proteins and peptides due to the large relative surface areas of these wells. Even moving this prep to 384 well plates will undoubtedly lead to greater peptide, and likely drug recovery. ^23^ The ability to quantify drugs in single human cells with nanoflow chromatography on one of the most sensitive high resolution mass spectrometers ever designed ^35^ also seems less impressive given the recent successes in lipid analysis in single cells using last generation instruments at analytical flow rates. ^36^ This ultimately suggests that the measurement of drug concentrations and simultaneous sampling of proteomics, and possibly other small molecules, may be feasible on other hardware. Finally, the processing and analysis of these data can be easily performed with the same proteomic and small molecule analysis software the mass spectrometry field knows and uses. The greedy dream of analyzing multiple classes of molecules within a single cell isn’t new, Fulcher *et al*., recently described the partitioning of protein from transcripts and the analysis of both from single cells ^22^ because a comprehensive approach is what will be necessary to solve the toughest biological riddles.

## Acknowledgements

I’d like to thank Megan Rigby, Alexis Norris and James Fulcher for conversations and insight critical to this work.

